# In the spotlight, losing IR region: insights from euglenophytes challenge the importance of inverted repeats in the genomes of secondary plastids

**DOI:** 10.1101/2022.08.04.502791

**Authors:** Kacper Maciszewski, Alicja Fells, Anna Karnkowska

## Abstract

Plastids, similarly to mitochondria, are organelles of endosymbiotic origin, which retained their vestigial genomes (ptDNA). Their unique architecture, commonly referred to as the quadripartite (four-part) structure, is considered to be strictly conserved; however, the bulk of our knowledge on their variability and evolutionary transformations comes from studies of the primary plastids of green algae and land plants. To broaden our perspective, we obtained seven new ptDNA sequences from freshwater species of photosynthetic euglenids – a group which obtained secondary plastids, known to have dynamically evolving genome structure, via endosymbiosis with a green alga. Our analyses have demonstrated that the evolutionary history of euglenid plastid genome structure is exceptionally convoluted, riddled with losses and multiple subsequent regains of inverted ribosomal operon (rDNA) repeats, as well as independent acquisitions of tandemly repeated rDNA copies. Moreover, we have shown that inverted repeats in euglenid ptDNA do not share their genome stabilizing property documented in chlorophytes. We hypothesize that the degeneration of the quadripartite structure of euglenid plastid genomes is connected to the group II intron expansion. These findings challenge the current global paradigms of plastid genome architecture evolution, and underscore the often-underestimated divergence between the functionality of shared traits in primary and complex plastid organelles.

## 1. Introduction

Approximately 1.5 billion years ago, in the Proterozoic eon, eukaryotes acquired the ability of photosynthesis through endosymbiosis between a heterotrophic host and a photosynthetic cyanobacterial cell, which gave rise to primary plastids (Archibald, 2015). This event changed life on Earth forever, as the ancient archaeplastid ancestor radiated into the diverse plastid-bearing lineages – land plants, red and green algae, which subsequently spread the photosynthetic capabilities to other groups of eukaryotes via secondary endosymbioses (Howe et al., 2008; Keeling, 2010). The integration of the cyanobacterial cells (and, later, primary plastid-bearing eukaryotic cells) with their new hosts involved massive endosymbiotic gene transfer from the symbiont’s genome into the host nucleus, leading to extreme streamlining of the plastid genome (plastome, ptDNA) (Burki et al., 2014; Ponce-Toledo et al., 2019).

However reduced, plastid genomes are retained in rather similar form across plastid-bearing lineages – they are almost invariably single, circular molecules, ranging from 50 to 200 kbp in size, carrying a vestigial gene repertoire encompassing predominantly photosynthesis-related genes and a major part of host-independent gene expression apparatus (de Vries and Archibald, 2018, 2017; Maier and Schmitz-Linneweber, 2004). What is more, ptDNA organization is also conserved to some extent, with the quadripartite structure – comprising a small (SSC) and large (LSC) single-copy region, flanked by a pair of inverted repeats (IRs) – being the most typical (Palmer and Thompson, 1982; Turmel et al., 2017, 2015; Zhu et al., 2016).

Still, as our state of knowledge on plastomes of the secondary plastid-bearing lineages expanded, it became evident that the quadripartite structure is strictly conserved mostly in land plants and green algae, while in others, the ptDNA architecture is substantially more diverse (Han et al., 2019; Kamikawa et al., 2015a; Karnkowska et al., 2018; Oborník and Lukeš, 2015; Turmel et al., 2017, 2015; Zhu et al., 2016), including some truly spectacular outliers, such as linear, split into minicircles, or even branching forms (Smith and Keeling, 2015). As a note, plastid IRs constitute a very particular case of a vast category of prokaryotic genetic elements collectively referred to as inverted repeats, as they contain the ribosomal subunit genes (*rrn16, rrn23, rrn5*), as well as transfer RNA genes and, in many taxa, protein-coding genes. In the following work, the phrase “inverted repeats” will always refer to the plastid-exclusive type of this structure (Lavi et al., 2018; Turmel et al., 2017).

The current theory states that the role of the inverted repeats in plastomes is mainly genome stabilization – they constitute a part of machinery for DNA repair via homologous recombination, which, in turn, is proposed to be responsible for both the lower substitution rate and less frequent genome rearrangements in IR-bearing plastid genomes in comparison with IR-deficient ones (Jin et al., 2020; Maréchal and Brisson, 2010; Palmer and Thompson, 1982; Turmel et al., 2017; Zhu et al., 2016). Nonetheless, studies of these phenomena have thus far been limited to the primary plastid-bearing taxa (e.g., land plants and chlorophytes), while others, even despite the abundance of genomic data on these organisms, remain rather neglected.

A perfect example of a secondary plastid-bearing group of algae, constituting a showcase for ptDNA architecture diversity, are the photosynthetic euglenids (Euglenophyta). This rather small (comprising below 20 genera, divided into three families – Euglenaceae, Phacaceae and Eutreptiaceae) and relatively young (plastid acquisition is estimated to have occurred approximately in the Ediacaran period (Jackson et al., 2018)) group of algae has attracted researchers’ attention for centuries due to their immense abundance in the freshwater environments and captivating morphology (Kostygov et al., 2021; Marin et al., 2003; Novák Vanclová et al., 2020). What is more, the earliest studies of their plastid genomes revealed an array of unique hallmark traits, such as multiple tandemly-repeated copies of the ribosomal operon or explosive group II intron expansion, prompting further investigation, which has so far resulted in over 30 full or partial ptDNA sequences of euglenophytes having been published up to date. These, in turn, made it possible to describe other features of divergent evolution in this group, such as the horizontal acquisition of maturase genes, but also rather broad variability of plastid genome structure, including three main types of rDNA repeat organization: single copy, inverted repeats, and tandem repeats (Bennett et al., 2012; Karnkowska et al., 2018; Maciszewski et al., 2022; Wiegert et al., 2012).

As transitions between organization types and their evolutionary consequences in euglenid plastid genomes remain documented, but not deeply investigated, we aimed to focus on this aspect of plastome evolution in the following study. Thus, we have selected seven species of freshwater photosynthetic euglenids (Euglenales), whose positions are close to the known nodes on their phylogenetic tree on which ptDNA structure rearrangements have most likely occurred, and sequenced their plastid genomes in order to broaden our scope of investigation for euglenids as a model group for studying plastome evolution and, having combined the new data with the aforementioned substantial body of reference, to test the long-standing hypothesis on correlation between IR conservation and diminished mutation rate outside of the primary plastid-bearing organisms.

## 2. Materials and methods

### 2.1. Research subjects, cultivation and isolation of the genetic material

For the purpose of this work, we cultivated seven strains of photosynthetic freshwater euglenids: *Euglena agilis* ACOI 2790, *Euglena deses* CCAP 1224/20, *Euglena undulata* MI03, *Strombomonas costata* ACOI 2992, *Colacium mucronatum* SAG 1211-1, *Phacus warszewiczii* ASW08064, and *Flexiglena variabilis* Boża Wola strain. Optimal growth of the microorganisms was observed on liquid S2T2 medium prepared according to the recipe on the ACOI culture collection website (http://acoi.ci.uc.pt), supplemented with 4 μg/ml of vitamin B12 and a single autoclaved pea seed (*Pisum sativum*), maintained in a room temperature light cabinet with 16/8h light/dark cycle in 10 ml glass tubes. Satisfactory culture density was determined by microscopic observations, after which the cultures were centrifuged for 3 mins at 3000 rpm.

Total DNA isolation from cell pellets was performed using DNeasy Blood & Tissue Kit (QIAGEN, USA) according to the manufacturer’s protocol, including an additional step of RNA digestion using RNase A. Quality control of the obtained isolates was carried out via spectrophotometric analysis using an Implen NP80 NanoPhotometer (Implen GmbH, Germany).

### 2.2. High-throughput DNA sequencing and plastid genome assembly

Total DNA samples of the seven euglenid strains were handed to an external company (Genomed S.A., Warsaw, Poland) for high-throughput sequencing using Illumina MiSeq platform. The sequencing yielded paired-end reads of different length, depending on the library preparation method used: 250 bp for *E. agilis* (approximately 7.2 million reads), *E. deses* (4.4 million reads), *P. warszewiczii* (8.2 million reads) and *S. costata* (4.0 million reads), and 300 bp for *C. mucronatum* (5.6 million reads), *E. undulata* (4.2 million reads) and *F. variabilis* (3.5 million reads). Quality control of the sequencing libraries was carried out using FastQC v0.11.6 tool (Andrews, 2010), and data trimming (i.e., removal of the Illumina Universal Adapter sequences) was performed using Trimmomatic v0.39 tool (Bolger et al., 2014).

Initial genome assembly of all datasets was performed using SPAdes v3.15.2 (Prjibelski et al., 2020), followed by identification of plastid genome fragments among the assembled contigs via BLASTN algorithm (Altschul et al., 1990) using publicly available euglenid ptDNA sequences as queries. Largest contigs identified as plastid genome fragments were extracted and used as seed sequences for plastid genome assembly using NOVOPlasty v4.3.1 (Dierckxsens et al., 2017). Although all plastid genomes have been successfully circularized, additional quality control was employed to verify their completeness: all plastid genome hits were extracted and visualized in Bandage v0.8.1 software (Wick et al., 2015) to confirm the circularization of the assembly; additionally, raw reads were mapped onto the NOVOPlasty assemblies using Bowtie v2.2.6 (Langmead and Salzberg, 2012) and Samtools v1.6 (Li et al., 2009) and the coverage per nucleotide position was calculated using Bedtools v2.25.0 (Quinlan and Hall, 2010) to detect putative low-coverage regions which would indicate misassembly.

### 2.3. Plastid genome annotation and visualization

Annotation of the obtained ptDNA sequences was carried out in Geneious Prime v2022.1.1 software (https://www.geneious.com) using Live Annotate & Predict toolkit (Find ORFs and Annotate From… features), utilizing a manually constructed database of all published euglenid plastid genomes as reference data for gene annotations. Identities and exon boundaries of all protein-coding genes were confirmed by cross-referencing with the NCBI non-redundant protein database (NCBI-nr) via BLASTX algorithm (Altschul et al., 1990) and the PFAM 35.0 protein families’ database (pfam.xfam.org) using the browser-accessible internal HMM search feature (Mistry et al., 2021). Plastid genome maps were generated using the OGDraw v1.3.1 online tool (Greiner et al., 2019).

### 2.4. Plastid genome-based phylogenomic analysis

58 protein-coding genes of non-ambiguous origins and function (i.e., excluding intron maturase genes, such as *roaA, ycf13* or *mat2/4/5/6/7*) were extracted from the annotated ptDNA sequences, translated to amino acid sequences, and combined with their homologues from 32 published plastid genome sequences of Euglenophyta and four published plastid genome sequences of Pyramimonadales (Chlorophyta). Protein sequences were aligned using L-INS-i method in MAFFT v7.310 (Katoh and Standley, 2013), and the single-gene alignments were concatenated using catsequences script (https://github.com/ChrisCreevey/catsequences) to produce a data matrix with total length of 18,143 amino acids.

The concatenated alignment was used as the input for phylogenetic analyses via maximum likelihood method implemented in IQ-TREE v2.0.6 software (Minh et al., 2020), and via Bayesian inference method implemented in MrBayes v3.2.6 (Ronquist et al., 2012). Maximum likelihood phylogeny reconstruction used a partitioned matrix with automatic substitution model selection for each partition (*-m TEST* parameter), and 1000 non-parametric bootstrap replicates. Bayesian reconstruction used a non-partitioned dataset with preset sequence evolution model (cpREV), with 1 000 000 generations (incl. 250 000 generations burn-in), after which convergence of the Markov chains was achieved. Both methods yielded fully congruent tree topology.

### 2.5. Ancestral state reconstruction

Reconstruction of ancestral states of plastid genome organization (IR-bearing versus IR-less) was carried out using corHMM v2.7 package (Beaulieu et al., 2013) in R v4.1.3, implemented in R Studio 2022.02.0 Build 443 (RStudio Team, 2020). IR presence was encoded as a binary trait, mapped (and plotted via plotRECON command) onto the tree topology obtained via Bayesian reconstruction. Four manually constructed substitution matrices were tested: with equal rate of state transition, with unequal rate of state transition, with only 0->1 transition possible, and with only 1->0 transition possible.

### 2.6. Rate of evolution estimation

For protein-coding genes, codon alignments for all single gene clusters were prepared using PAL2NAL v14 software (Suyama et al., 2006). Rates of synonymous and non-synonymous substitutions (*dN/dS*) for all gene alignments were calculated using CodeML tool implemented in PamlX v1.3.1 toolkit (Xu and Yang, 2013). Mean *dN/dS* values were calculated for two groups of euglenophytes: IR-bearing (13 taxa) and IR-less (26 taxa) for all 58 genes, and compared using two-sided Mann-Whitney *U*-test implemented in an online Social Science Statistics calculator (https://www.socscistatistics.com/tests/mannwhitney/).

For rRNA genes, nucleotide sequence alignments were prepared using L-INS-i method in MAFFT v7.310 (Katoh and Standley, 2013), and the two alignments (*rrn16, rrn23*) were concatenated using catsequences script (https://github.com/ChrisCreevey/catsequences) to produce a data matrix with total length of 5,954 nucleotides. The concatenated alignment was used as the input for phylogenetic analysis via maximum likelihood method implemented in IQ-TREE v2.0.6 software (Minh et al., 2020) with automatic substitution model selection (*-m TEST* parameter), and 1000 non-parametric bootstrap replicates, in two variants: with no constraints, and with constrained topology based on the results of the plastid protein-coding genes-based phylogeny. For both phylogenies, mean branch length values were calculated for branches divided into two groups: reconstructed as IR-bearing (19 branches in total), and reconstructed as IR-less (52 branches in total). Branches at which state transitions occurred were not taken into account. Mean values were subsequently compared using two-sided Mann-Whitney *U*-test implemented in an online Social Science Statistics calculator (https://www.socscistatistics.com/tests/mannwhitney/).

## 3. Results and discussion

### 3.1. Plastid genome characteristics and phylogeny

The basic characteristics of the seven new ptDNA sequences of freshwater euglenophytes have been shown in Table 1, and their structure has been depicted on Supplementary Figure 1. As expected, based on the past studies (Bennett et al., 2012; Karnkowska et al., 2018; Maciszewski et al., 2022), the sequenced euglenid plastid genomes do not exhibit vast diversity of genetic repertoire – ranging from 88 genes in *E. undulata* to 101 in *S. costata* – with the variable numbers of rDNA operon copies and group II intron maturase genes accounting for almost all of the differences in gene content between the investigated taxa.

**Table 1.** Characteristics of the seven novel plastid genomes of Euglenales presented in this study. Numbers and length of introns do not include twintrons.

|  | *Colacium mucronatum* | *Euglena agilis* | *Euglena deses* | *Euglena undulata* | *Flexiglena variabilis* | *Phacus warszewiczii* | *Strombomonas costata* |
| --- | --- | --- | --- | --- | --- | --- | --- |
| Length | 147,974 bp | 107,929 bp | 83,748 bp | 92,487 bp | 105,410 bp | 84,299 bp | 185,621 bp |
| GC content | 25.6% | 26.5% | 26.5% | 27.3% | 27.3% | 25.6% | 27.5% |
| Total no. of genes | 94 | 93 | 94 | 88 | 94 | 95 | 101 |
| Protein-coding genes | 60 | 62 | 61 | 58 | 63 | 63 | 68 |
| tRNA genes | 29 | 28 | 27 | 28 | 28 | 29 | 27 |
| rRNA genes | 5 | 3 | 6 | 2 | 3 | 3 | 6 |
| Introns | 119 | 97 | 72 | 78 | 83 | 63 | 129 |
| Total intron length | 62,991 bp | 40,630 bp | 21,627 bp | 29,591 bp | 46,955 bp | 23,347 bp | 115,766 bp |

In contrast, the investigated strains displayed substantial differences in total ptDNA size, ranging from approximately 83.7 kb in *E. deses* to over twice that size, 185.6 kb, in *S. costata*. What is more, plastomes of *S. costata* and *C. mucronatum*, both examined in our study, are currently the two largest among all euglenophytes, with the latter (147.9 kb) also exceeding the size of all previously published plastid genomes of this group. However, the numbers of functional genes (i.e., protein-coding genes as well as tRNA and rRNA genes) are not even a noticeable factor influencing total ptDNA size in euglenophytes – although the plastid genome of *S. costata* is indeed the most gene-rich among the seven new ptDNA sequences, no such trend was visible among other examined plastomes. Instead, the total plastid genomes size differences can be almost entirely attributed to non-coding sequences – in particular, the group II introns, whose explosive expansion is among the most distinctive traits of euglenid plastids (Karnkowska et al., 2018; Maciszewski et al., 2022).

The plastid genome-based phylogeny of euglenophytes, obtained in our study (Figure 1), is almost fully congruent with the most recent nuclear and plastid rDNA-based reconstructions (Karnkowska et al., 2018; Kim et al., 2015; Maciszewski et al., 2022). Only one minor discrepancy was observed: in our reconstruction, *E. longa* is a sister taxon to a clade comprising *E. gracilis* and *E. hiemalis*, instead of being sister to only *E. hiemalis*. This, however, does not impact the results of our further analyses, as they relate to clades which possess identical values for traits studied in our work. Combined with the predominantly absolute or very high bootstrap support and posterior probability values for the plastid-based phylogeny presented here, as well as fully congruent topology between Bayesian and maximum-likelihood reconstructions in this study and the previous works (Karnkowska et al., 2018, 2014; Linton et al., 2010; Maciszewski et al., 2022), it is reasonable to assume that the euglenophyte phylogeny shown on Figure 1 is the most stable and credible one up to date. Nonetheless, certain discrepancies between phylogenies based on molecular markers of diverse origin (nuclear versus organellar), sequence type (nucleotide versus protein) and alignment size (single genes versus concatenated multigene matrix) are to be expected.

**Figure 1.**
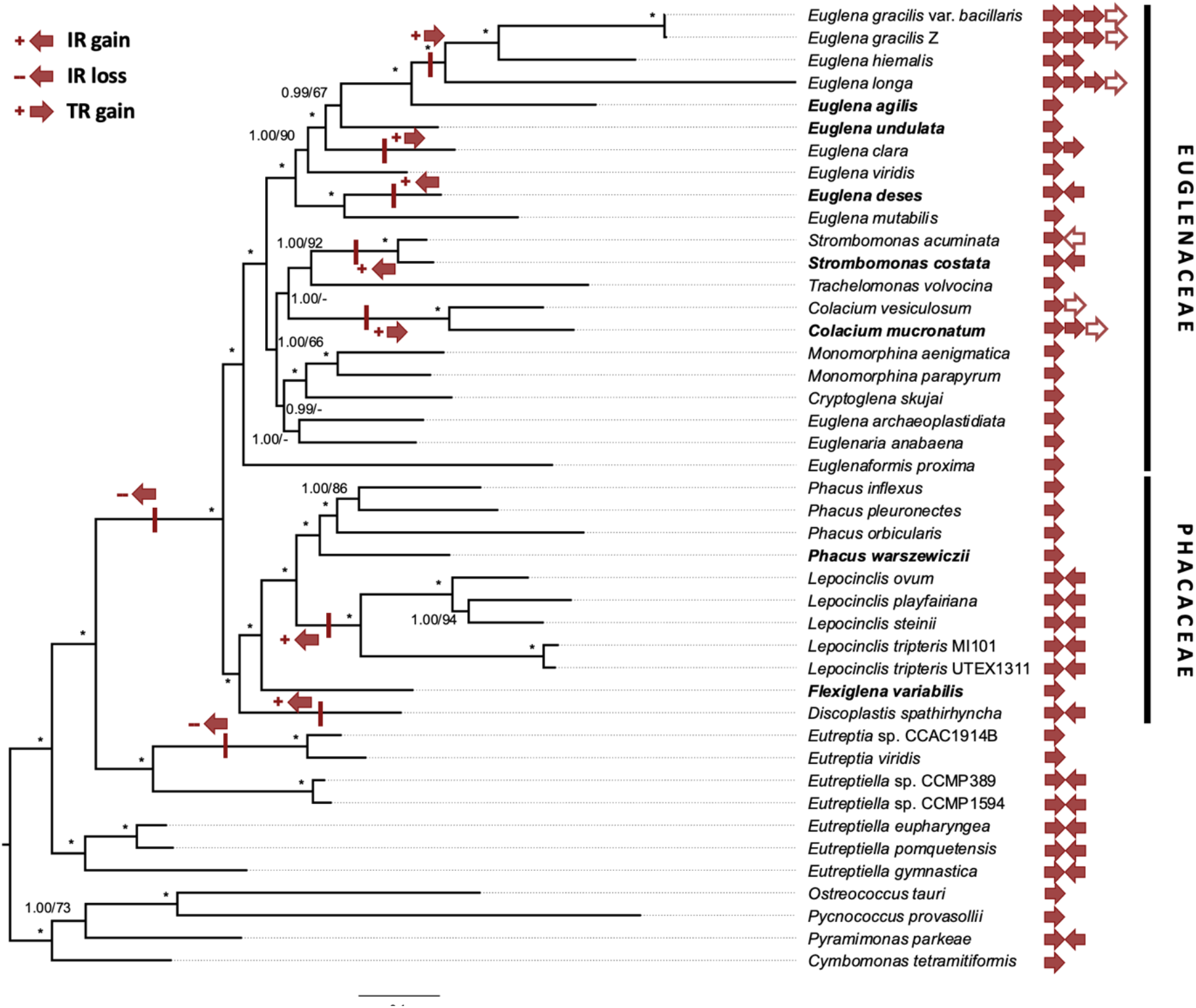
Plastid-based phylogenomic tree of Euglenophyta. Species names in **bold** indicate organisms whose ptDNA was first sequenced in this work. rDNA operon copy number (filled arrows denote complete operon copies; hollow ones denote incomplete ones) and orientation are shown on the tree tips; gains and losses are marked on the tree branches. Bayesian posterior probability and bootstrap support values above 50 are shown at the nodes. Asterisks (*) denote absolute probability and support (>0.99/>95).

### 3.2. Unexpected gains and losses of the rDNA copies

As demonstrated in past studies, ptDNA organization in euglenophytes has undergone substantial diversification over time, encompassing not only the group II intron expansion, but also gains and losses of rDNA copies – the most recent reconstruction suggests three independent losses of one of the inverted repeats: in the ancestors of genera *Eutreptia* and *Phacus*, and in the common ancestor of the family Euglenaceae (Karnkowska et al., 2018). Moreover, certain species of the genus *Euglena* have acquired additional copies of the ribosomal operon, situated consecutively in the same orientation in the genome, resulting in a unique genetic structure, commonly referred to as tandem repeats (Gockel and Hachtel, 2000; Hallick et al., 1993; Hewadikaramge and Linton, 2018). However, the organization of the seven new euglenid plastid genomes, when mapped onto the studied group’s phylogeny (Figure 1), challenges nearly all assumptions of the previously proposed model of three IR losses and a single TR gain.

First of all, plastid genomes of *E. agilis* and *E. undulata* – both of which are situated within a clade of *Euglena* spp. previously proposed to possess TRs (see Figure 1) – carry a single rDNA copy, indicating either two independent gains of a TR within the genus *Euglena*, or alternatively (and less parsimoniously), a gain and two independent losses. Moreover, we identified yet another independent gain of tandem repeats, this time outside of the genus *Euglena*, specifically: in *C. mucronatum*, which strongly indicates that single-copy rDNAs in euglenid plastids are quite likely to undergo duplication, forming TRs. This assumption is also supported by the observation that the TR copy number is also varied among *Euglena* and *Colacium* spp. – *E. clara* and *E. hiemalis* possess two full copies, while *C. mucronatum* possesses “two and a half” (two full copies and an additional *rrn16* gene), and *E. gracilis* and *E. longa* possess “three and a half” copies (three full copies and an additional *rrn16* gene). It is also worth mentioning that our study is not the first to obtain a ptDNA sequence of a representative of the genus *Colacium;* however, the published sequence from *C. vesiculosum* is incomplete, with the missing part most likely including a part of a ribosomal operon repeat, which would make any conclusions on IR/TR evolution based on that sequence at least dubious (Wiegert et al., 2013).

Secondly, *E. deses* and *S. costata* – both representing Euglenaceae, which were proposed to have ancestrally lost the inverted rDNA repeats – do, in fact, possess IRs. This finding is particularly puzzling because of the position of these two species on the phylogenetic tree, indicating that, for the observed IR loss and retention pattern to appear, there must have been six independent IR losses within the Euglenaceae alone: in the ancestor of *E. gracilis/hiemalis/longa/agilis/undulata/clara/viridis*, in *E. mutabilis*, in the genus *Trachelomonas*, in the genus *Colacium*, in the genus *Euglenaformis*, and in the ancestor of the genera *Monomorphina, Cryptoglena* and *Euglenaria*. Bearing this in mind, the ancestral IR loss in Euglenaceae (as proposed by Karnkowska et al., 2018; see Figure 1), followed by two independent regains in *E. deses* and *S. costata*, would be a substantially more parsimonious explanation for the pattern observed in the extant species.

Last, but not least, we found *F. variabilis* – a representative of the newest described euglenophyte genus, *Flexiglena* (Łukomska-Kowalczyk et al., 2021) – to possess a single rDNA copy, even though its phylogenetic position among predominantly IR-bearing taxa (*Lepocinclis* and *Discoplastis* spp.; see Figure 1) suggested that it is rather likely to carry IRs as well. In contrast with the other freshwater euglenophyte family, in Phacaceae the observed IR loss and retention pattern has two comparably likely explanations: either two independent IR losses in *Phacus* and *Flexiglena* and their retention in *Lepocinclis* and *Discoplastis*, or two independent gains in *Lepocinclis* and *Discoplastis* and retention of the ancestrally IR-less state in *Phacus* and *Flexiglena*. However, assuming that Euglenaceae were ancestrally IR-less as outlined above, a single loss in the ancestor of freshwater euglenophytes (Euglenales – Euglenaceae + Phacaceae; see the systematics in Kostygov et al., 2021) and subsequent regains (twice in Euglenaceae and twice in Phacaceae) would seem to be a more plausible model for the evolutionary path of the rDNA operon copies in the investigated group.

### 3.3. The original ptDNA organization in the euglenophyte ancestor and the revised history of transitions

To unravel the convoluted history of ribosomal operon organization in euglenid plastids, we performed computational reconstruction of the ancestral states on the investigated group’s phylogeny (Figure 2). Among the four tested models for transition rates between IR presence and absence, the model with unequal transition rate has been selected as best-fitting for the dataset with Akaike information criterion (AIC) value of 45.04, closely followed by a model with equal transition rate (AIC = 46.91), while unidirectional transition models were significantly worse (both with AIC = 2×10^5^). The reconstruction produced no ambiguous ancestral states on any node on the tree, with the IR presence in the ancestor of all euglenid plastids reconstructed at >99% probability.

**Figure 2.**
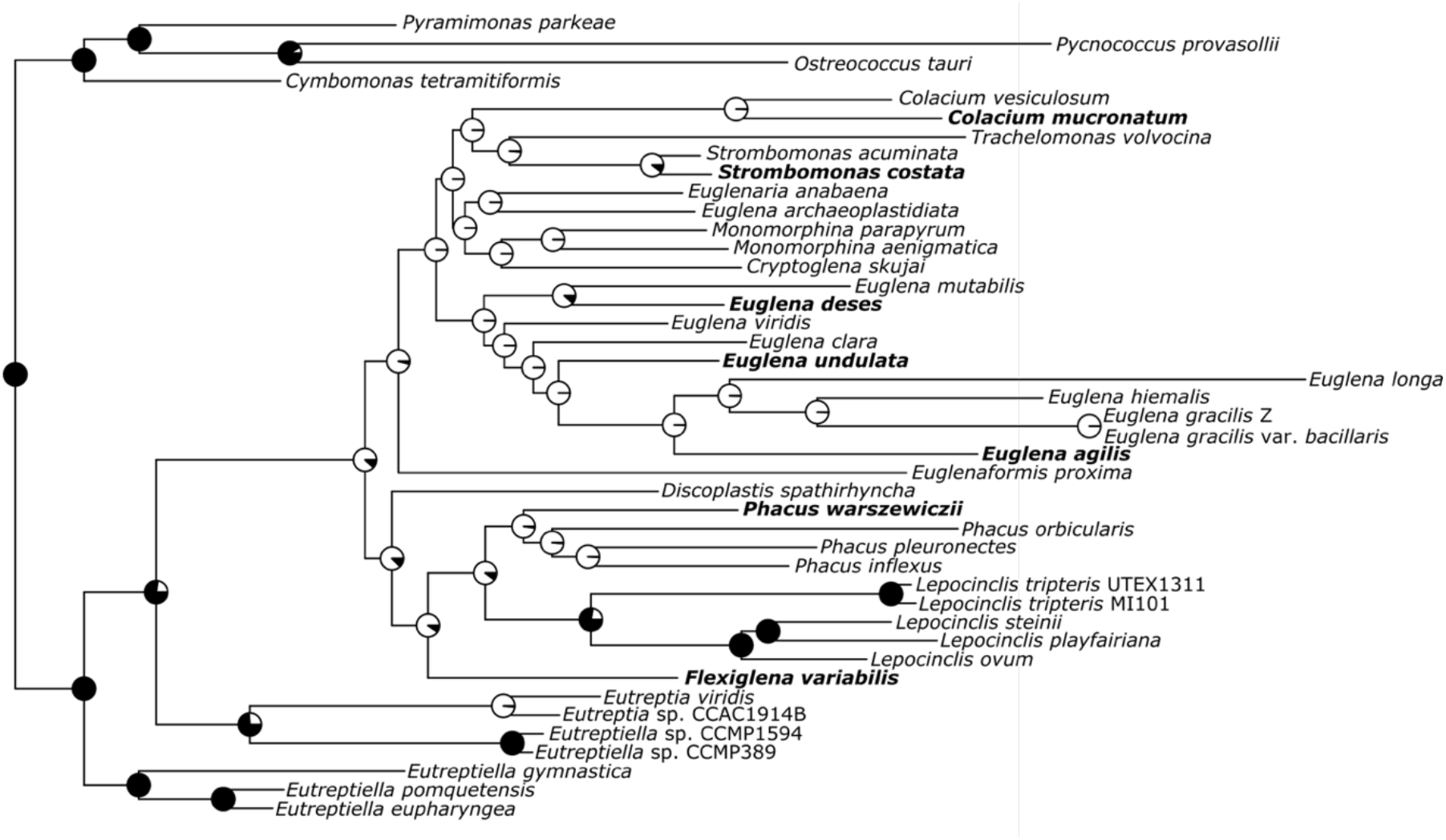
Ancestral state reconstruction of plastid genome organization in Euglenophyta, mapped onto the group’s phylogeny (see Figure 1). State transition rate was automatically determined as unequal, and this information has been included in the preset transition rate matrix. Hollow circles (○) at the nodes denote IR-less states; full circles (●) denote IR presence.

Moreover, the reconstructed states at the other nodes point toward the hypothesis outlined before: that the IRs were lost in euglenophytes only twice – in the ancestor of all freshwater euglenids (Euglenales), and in the ancestor of the genus *Eutreptia* (see Figure 2) – and were subsequently regained independently by *E. deses, S. costata, D. spathirhyncha* (or the genus *Discoplastis* – however, the lack of ptDNA sequences of its other species makes it impossible to determine), and the genus *Lepocinclis*. This stands in opposition to the hypothesis that IR gains, in contrast to losses, are very rare, and the only piece of strong evidence for such an occurrence ever taking place comes from the green algal genus *Chamaetrichon*, whose peculiar plastid genome possesses three IRs (Turmel et al., 2017, 2015).

On the other hand, the inverted repeats of euglenids are substantially different from their counterparts in primary plastids of plants or green algae. While the IRs in plastid genomes usually flank two single-copy regions containing most of the protein-coding genes and tRNA genes, those in euglenophytes are almost always adjacent to each other – the small single-copy region is very short and has no coding content, and the entire protein-coding gene repertoire is located in the large single-copy region (Karnkowska et al., 2018; Maciszewski et al., 2022). Only one exception has been documented so far – *Eutreptiella gymnastica*, where one of the repeats is split, and a six-gene insertion separates the *rrn16* and *rrn23* genes, thus constituting a unique kind of a small single-copy region (Hrdá et al., 2012). Nonetheless, despite the presence of IRs in some lineages, none of the euglenid plastid genomes actually carries a true quadripartite structure.

Additionally, all euglenid IRs have identical gene content of the ribosomal operon and two tRNA genes, in contrast with the substantially more complex inverted repeats known e.g. from green algae, which encompass a wide array of protein-coding genes (Turmel et al., 2017, 2015). Although it is quite difficult to state whether the euglenophyte plastids’ IRs are secondarily simplified, or that those in extant plants and algae gained their complexity late in their evolution and the simplicity is actually plesiomorphic, the adjacent position and small size of euglenid IRs makes it more plausible that they are products of numerous independent duplications.

### 3.4. Inverted repeat presence does not impact the rate of evolution of either protein-coding genes or rRNA genes

As the organization of euglenophyte ptDNA seems to have undergone numerous rearrangements over time, there does not seem to be strong evolutionary pressure for retention of any particular organization type. Previous studies have shown that the inverted repeats are conserved in plastid genomes due to their stabilizing activity as homologous recombination sites, as is evident from the substantially increased rate of sequence evolution and genomic rearrangements observed in IR-less taxa (Claude et al., 2022; Palmer and Thompson, 1982; Zhu et al., 2016). However, these analyses took only the data from primary plastids into account, leaving the diverse secondary plastid-bearing taxa understudied in this regard. Thus, given that there is a growing body of evidence for dissimilarity of evolutionary dynamics and selection intensity between primary and complex plastids (Uthanumallian et al., 2021), the missing piece of the puzzle may be tremendously important. To fill this information void, we have undertaken an analysis of the sequence evolution rate in ribosomal subunit genes, which form the bulk of the length of the plastid inverted repeats, as well as in protein-coding genes encoded outside of the IRs in the IR-bearing and IR-less plastomes of Euglenophyta.

We calculated *dN/dS* ratios based on the sequences of 58 plastid protein-coding genes in euglenid plastids, and obtained mean and standard deviation values for IR-bearing and IR-less taxa: *dN/dS* = 0.158 ± SD = 0.0378, and *dN/dS* = 0.197 ± SD = 0.0653, respectively. These values were compared using Mann-Whitney *U-*test, which yielded the *z-*score of -0.159 and the *p-*value of 0.436, which, quite interestingly, can be clearly interpreted as no difference between the investigated groups. Furthermore, our analysis of the rate of rDNA evolution – the small and large ribosomal subunit genes, which form the inverted repeats – produced congruent results in both studied variants of the rDNA-based phylogeny (Table 2), indicating no differences in the rate of evolution between the ribosomal subunit genes enclosed within IRs and ones present in single copies. These results together, being contradictory with analogical studies of green algae (Zhu et al., 2016) and land plants (Claude et al., 2022; Ping et al., 2021), constitute yet another key difference in the evolutionary paths of primary and secondary plastids and their genomes, pointing towards the loss of the stabilizing function of the IRs in this particular secondary plastid-bearing lineage.

**Table 2.** Comparison of the rates of evolution of rRNA genes in IR-bearing and IR-less euglenid plastid genomes estimated via phylogenetic tree branch length analysis.

|  | **Constrained phylogeny** | ***De novo*-reconstructed phylogeny** |
| --- | --- | --- |
|  | **Mean branch length ± SD** |  |
| **IR-bearing** | 0.0492 ± 0.0644 | 0.0473 ± 0.0604 |
| **IR-less** | 0.0543 ± 0.0577 | 0.0539 ± 0.0570 |
|  | **Mann-Whitney *U*-test's *z*-score; *p*-value** |  |
|  | 0.309; <u>0.757</u> | 0.127; <u>0.897</u> |

If the presence of the inverted repeats does not impact the rate of evolution of both protein-coding and rRNA genes in the analysed plastid genomes, it is only logical to assume that there is no discernible consequence to IR loss or retention in euglenophytes at all. We are therefore inclined to hypothesize that, contrary to primary plastid-bearing taxa, euglenid plastid genome organization follows the path of neutral evolution, with spontaneous rearrangements, such as rDNA copy gains or losses, being only retained as a result of the absence of the selective pressure to keep a specific genome architecture. Naturally, a conclusion that secondary plastids’ genome structure only evolves neutrally would be unsupported due to the lack of published results of analogical rate of evolution versus ribosomal operon organization analyses for taxa other than euglenids – to determine that, further studies are necessary.

Still, the euglenid plastid genome IRs remain a rather puzzling evolutionary peculiarity when a broader context is considered. Previous studies have demonstrated that inverted repeats are diverse genetic elements, ranging from several to thousands of nucleotides in length, spread across the entire tree of life, with multifaceted influence on the genome structure and evolution due to their capabilities for forming hairpins, and constituting the sites for flip-flop recombination and template switching during DNA replication. As a result, certain forms of IRs can have either generally stabilizing influence on the genome as constituents of DNA repair systems, as demonstrated in plant plastids, or, on the contrary, destabilizing impact as hotspots for mutations, as shown in bacteria and eukaryotic nuclei (Lavi et al., 2018; Maréchal and Brisson, 2010; Turmel et al., 2017). Therefore, it comes as a great surprise that in a certain genetic setting, IRs can have no noticeable impact on the genome’s evolution whatsoever. Moreover, taking into account that bacterial genomes can carry thousands of IR pairs, albeit very short, the curiosity lies not just in their retention in ptDNA, but in the fact that only a single pair is retained (Lavi et al., 2018).

It is also worth noting that IR losses in certain complex plastid-bearing algae, such as cryptophytes, have been proposed to be concomitant with loss of photosynthesis. This hypothesis is particularly interesting due to the presence of numerous parallels between euglenid and cryptophyte plastids, despite their different origins and vast evolutionary distance between the host lineages – e.g., independent group II intron expansion and acquisition, followed by partial degeneration, of intron-encoded maturase genes (Maciszewski et al., 2022; Suzuki et al., 2022). A connection between IR decay and a shift to heterotrophy is not likely in case of euglenids, since most of them lost IRs, while losses of photosynthesis in this lineage are comparably scarce. However, a possible link between intron accumulation and inverted repeat degeneration due to induced plastome instability and, additionally, the acquisition of new homologous recombination sites which turned IRs redundant, is certainly a plausible explanation for the cooccurrence of these two rather uncommon traits both in euglenid and cryptophyte plastids (Suzuki et al., 2022). Unfortunately, a solid proof for a link between group II intron expansion and IR redundancy is currently out of our reach, as it would require reference data from intron-less euglenid ptDNA, which have never been identified, while using data from pyramimonadalean green algae (the closest extant relatives of the euglenid plastid) could lead to erroneous conclusions due to the documented shift in the rate of plastid genome evolution following a endosymbiotic event (Uthanumallian et al., 2021).

## 4. Summary

In the presented work, we obtained full plastid genome sequences of seven species of freshwater photosynthetic euglenids, selected according to their phylogenetic positions within or adjacent to taxa known to have undergone ptDNA structure rearrangements in order to investigate the evolutionary dynamics of the genome organization within this model secondary plastid-bearing group. Only *Phacus warszewiczii*, simply shared the plastome structure of their closest relatives; however, we found many of the studied species to have divergent ribosomal operon organization – *Euglena agilis* and *Euglena undulata*, members of a TR-bearing clade, have a single rDNA copy, while *Colacium mucronatum*, located within a TR-less clade, has secondarily acquired tandem repeats. Similarly, *Euglena deses* and *Strombomonas costata*, both located within IR-less clades, possess IRs, while *Flexiglena variabilis*, branching within a predominantly IR-bearing clade, does not have them.

Our findings have demonstrated that the variability of ptDNA organization in euglenophytes, despite being studied before, is even more immense than previously thought.Our observation of the four independent secondary acquisitions of a rDNA inverted repeat not only challenges the hypothesis on the unlikelihood of formation of IRs *de novo*, but also impacts the more general theory of progressing reductive evolution of organellar genomes by constituting a prominent example of an increase, and not decrease, in an organellar genome’s complexity.

We therefore suggest that the reason behind the tremendous diversity in the architecture of the repeated ribosomal operon sequences lies in their partial loss of function: the secondary plastid of euglenophytes did not inherit the IR recombination-based repair mechanism, acting in the primary plastids of the Archaeplastida, and therefore the retention of IRs themselves offers little, if any, selective advantage. As a result, the photosynthetic organelle and its host do not bear any serious evolutionary consequences of gains and losses of additional copies of a short RNA-encoding genes, leading to formation of a multitude of divergent forms. Moreover, if other recombination sites, such as group II introns, have been introduced into the genome, the loss of redundant inverted repeats might even be beneficial.

Our understanding of the processes shaping the organellar genome structure and contents is, however, still limited. The next step worth taking to deepen it would be the broadening of the scope of the research subject by analyzing the genome structure and rates of evolution in other, especially more diverse complex plastid-bearing taxa, such as haptophytes or dinoflagellates. Subsequently, once more data is available, the plastid genome dynamics could be cross-referenced with the unique biological, physiological and genetic traits of different organisms with independently acquired plastids, which would help unravel the real significance of the evolutionary transformations investigated in this study. Furthermore, the intertwined influences of intron expansion, IR degeneration and photosynthesis losses would undoubtedly merit further investigation, and the mechanism which would compensate for the loss of the postulated key ptDNA repair system in other, intron-less plastid genomes still awaits discovery.

## 5. Acknowledgements

This work was supported by Preludium grant 2018/31/N/NZ8/01840 (National Science Centre, Poland) to KM, Excellence Initiative – Research University (IDUB) grant (Ministry of Science and Higher Education, Poland) to AF, and EMBO Installation Grant 4150 (to AK) and Ministry of Education and Science, Poland. We would like to sincerely thank our colleagues from the Institute of Evolutionary Biology (University of Warsaw, Poland): Jakub Baczyński, for his invaluable support in ancestral state reconstruction analyses, and Bożena Zakryś, for supporting us with her great experience and expertise in maintenance of algal cultures.

## 6. Author contributions

KM and AK conceptualized the study, interpreted the results and prepared the final version of the manuscript; AF assembled and annotated the plastid genome of *Flexiglena variabilis;* KM obtained the funding, formulated the hypotheses, assembled and annotated the other six plastid genomes, performed the bioinformatic analyses, and prepared the draft version of the manuscript with figures and tables; AK supervised the work.

## Appendix A. Supplementary figures

**Supplementary Figure 1.**
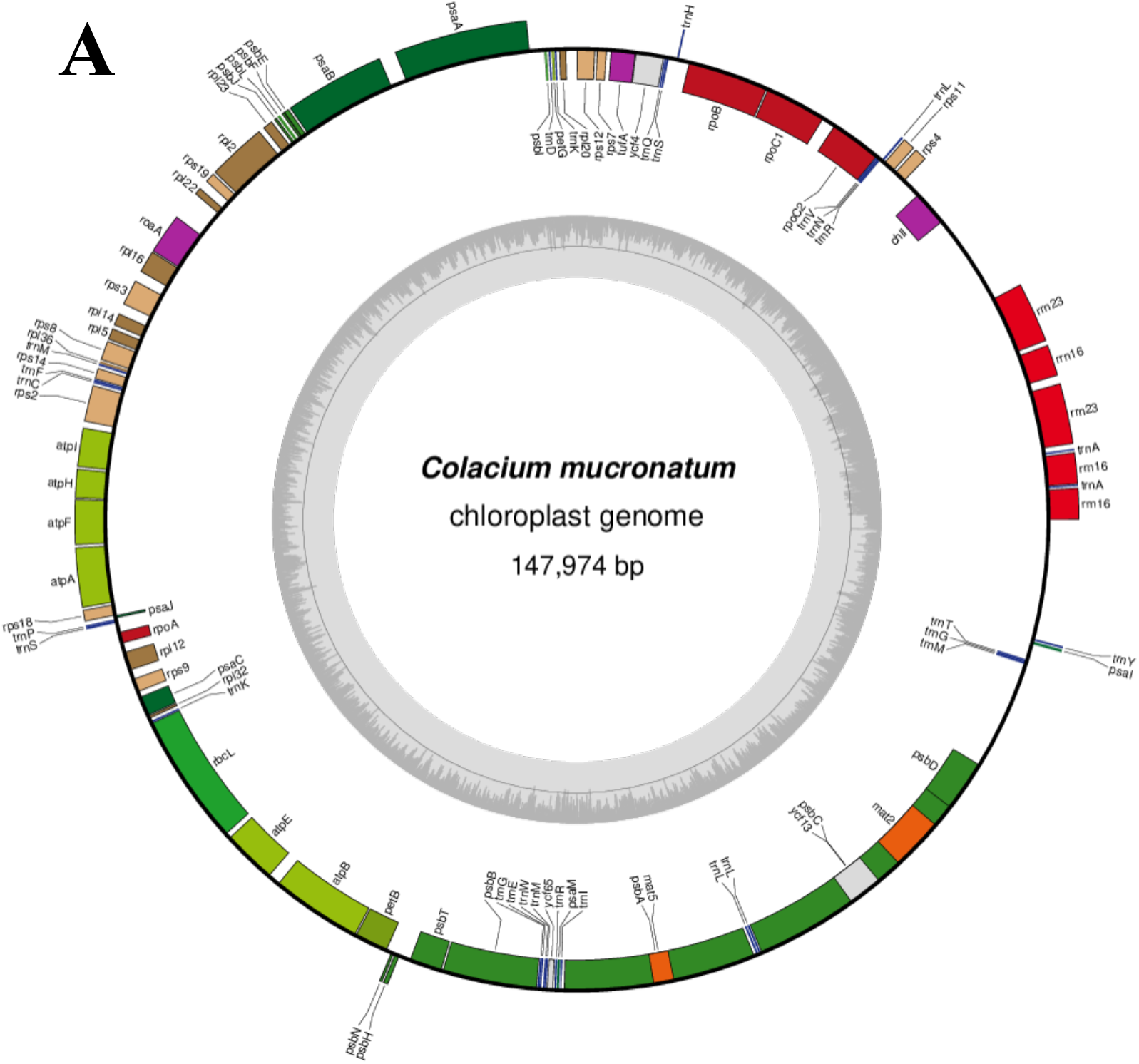
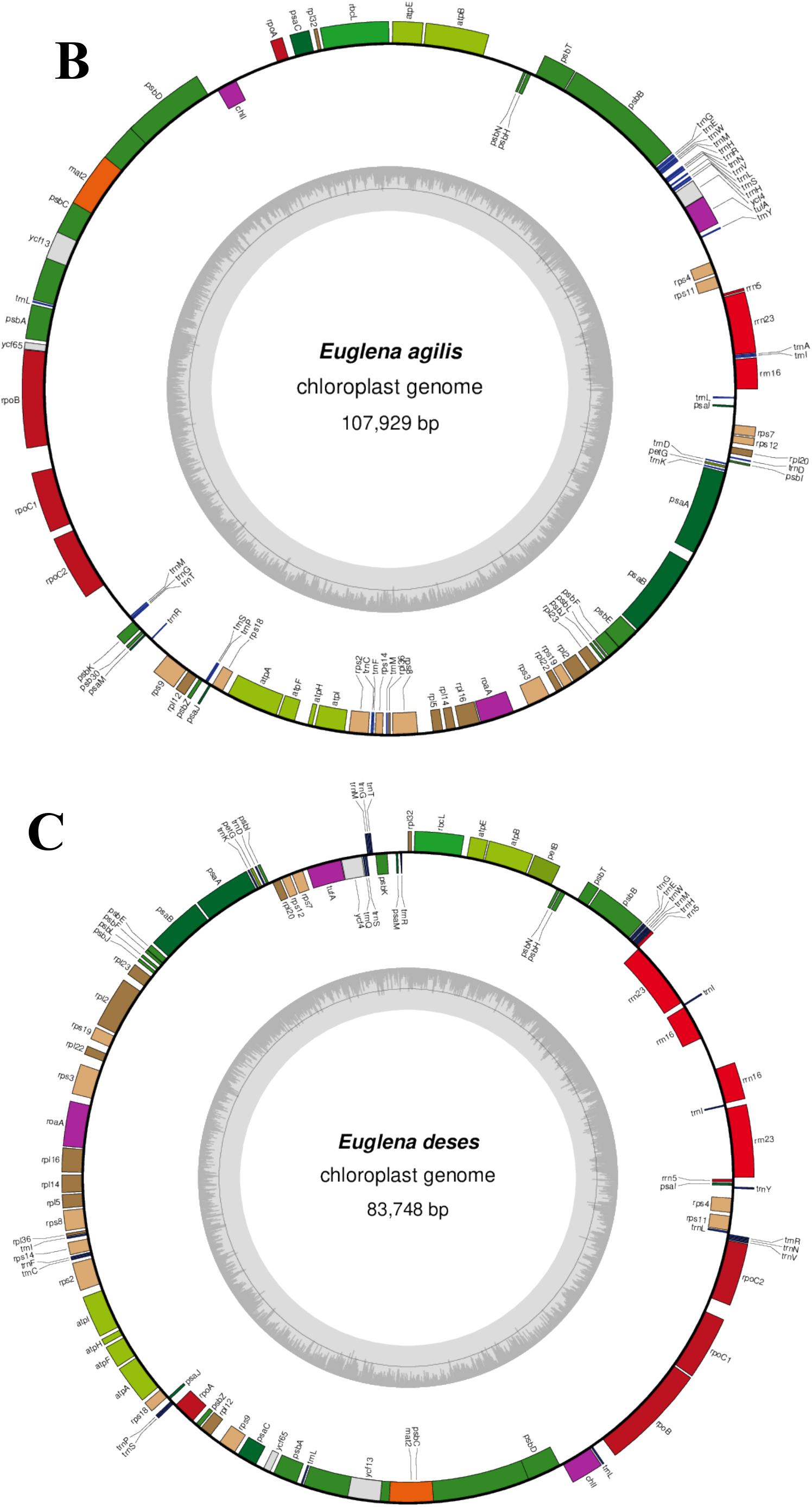
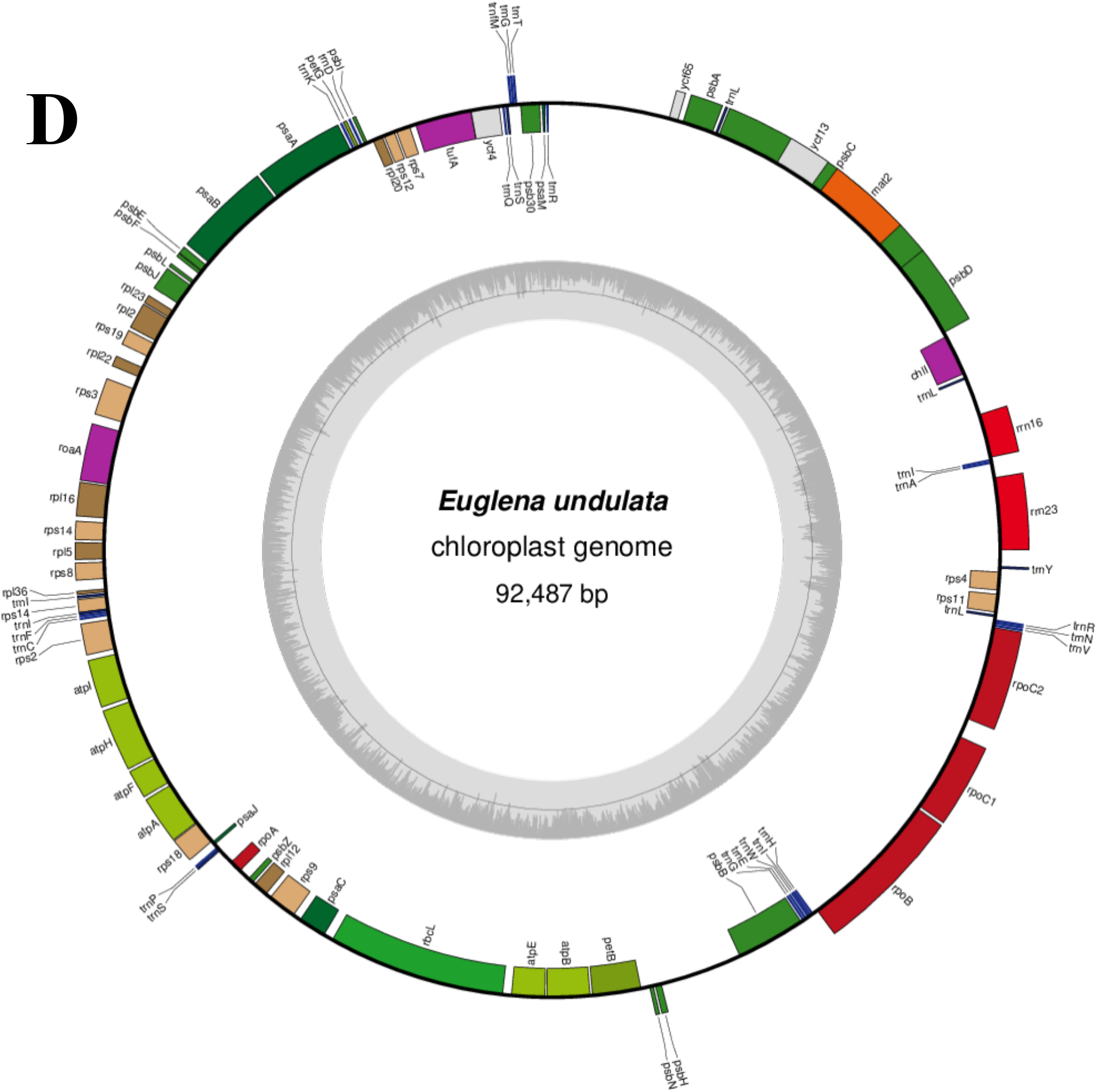
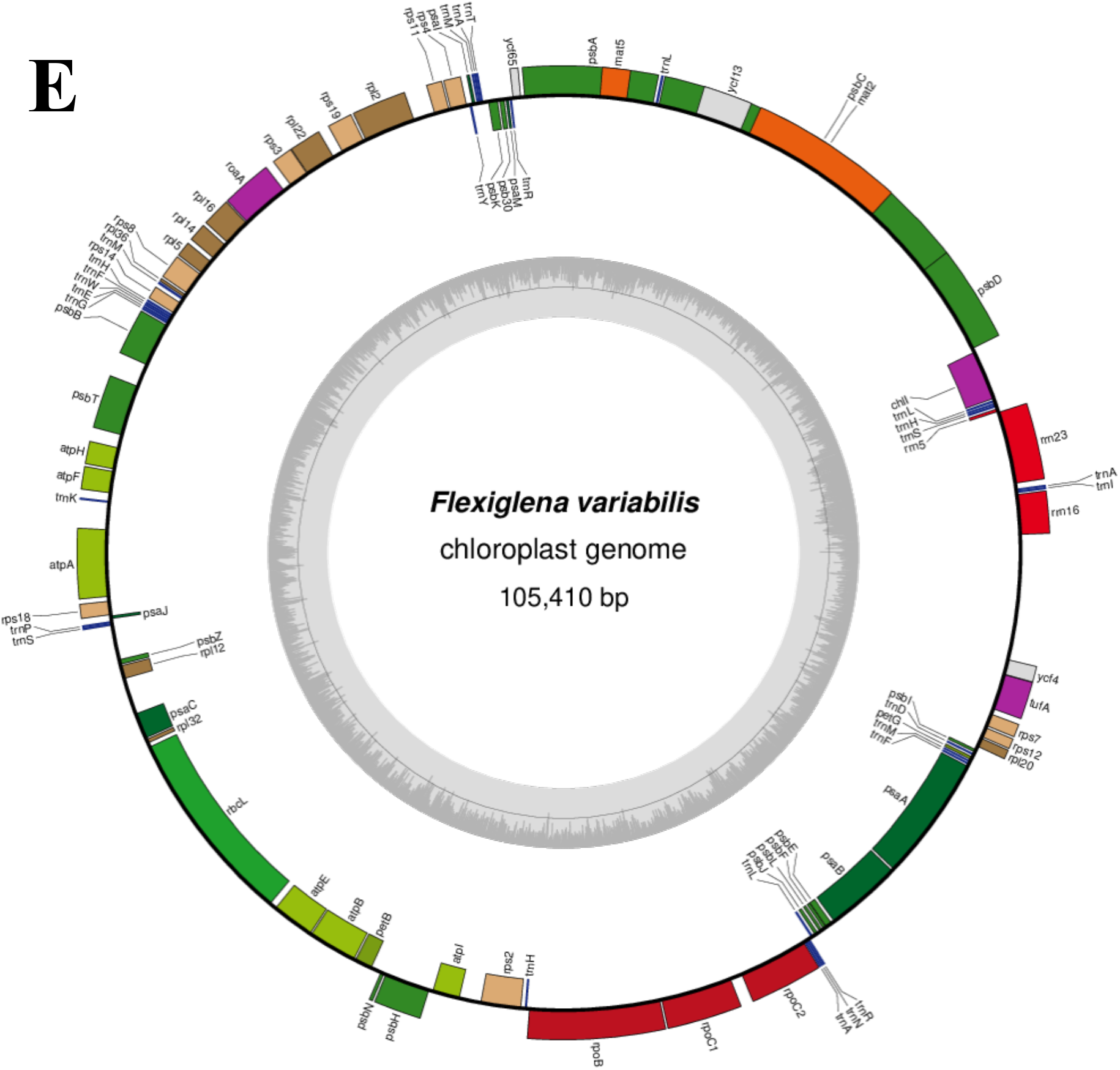
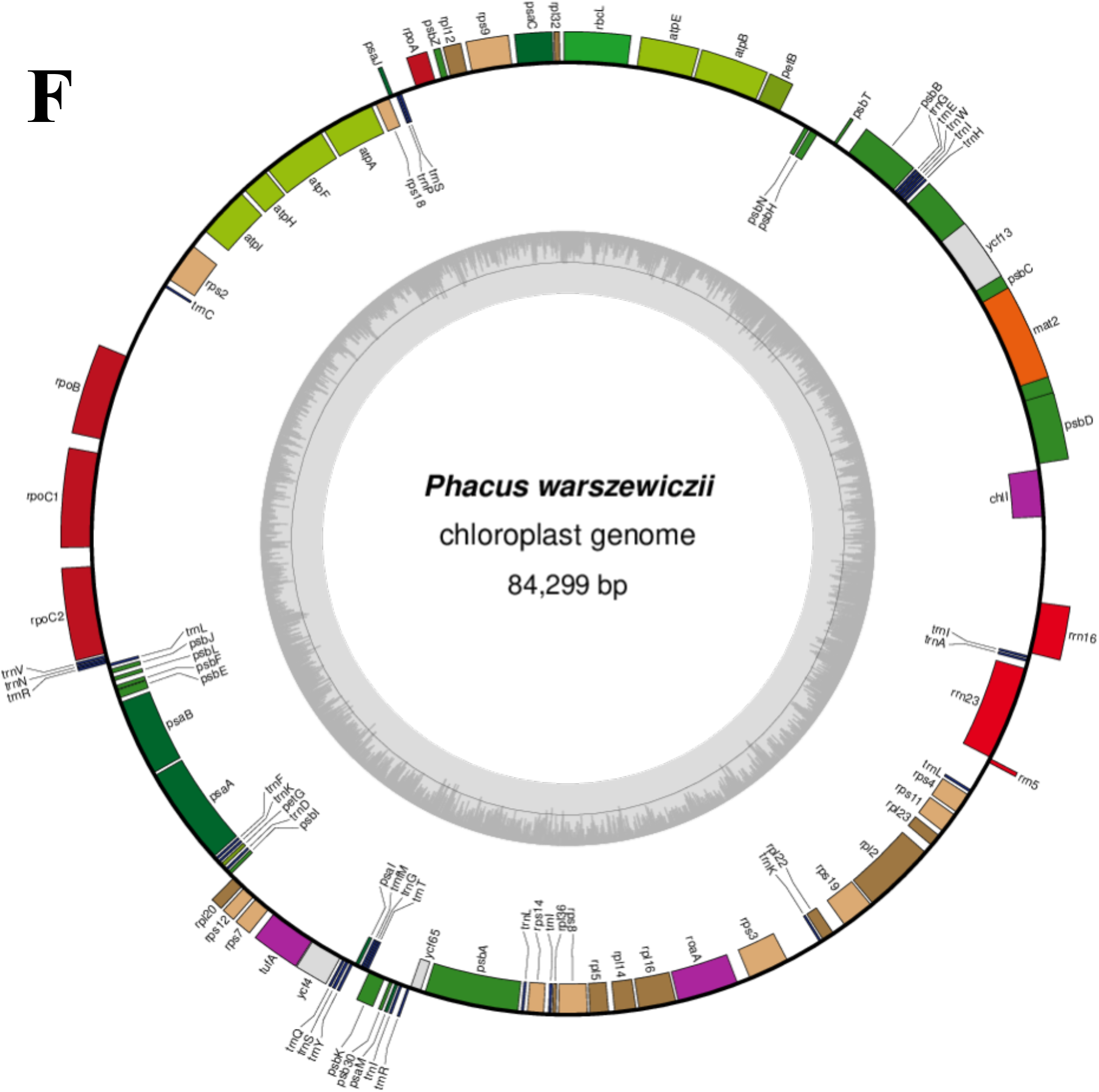
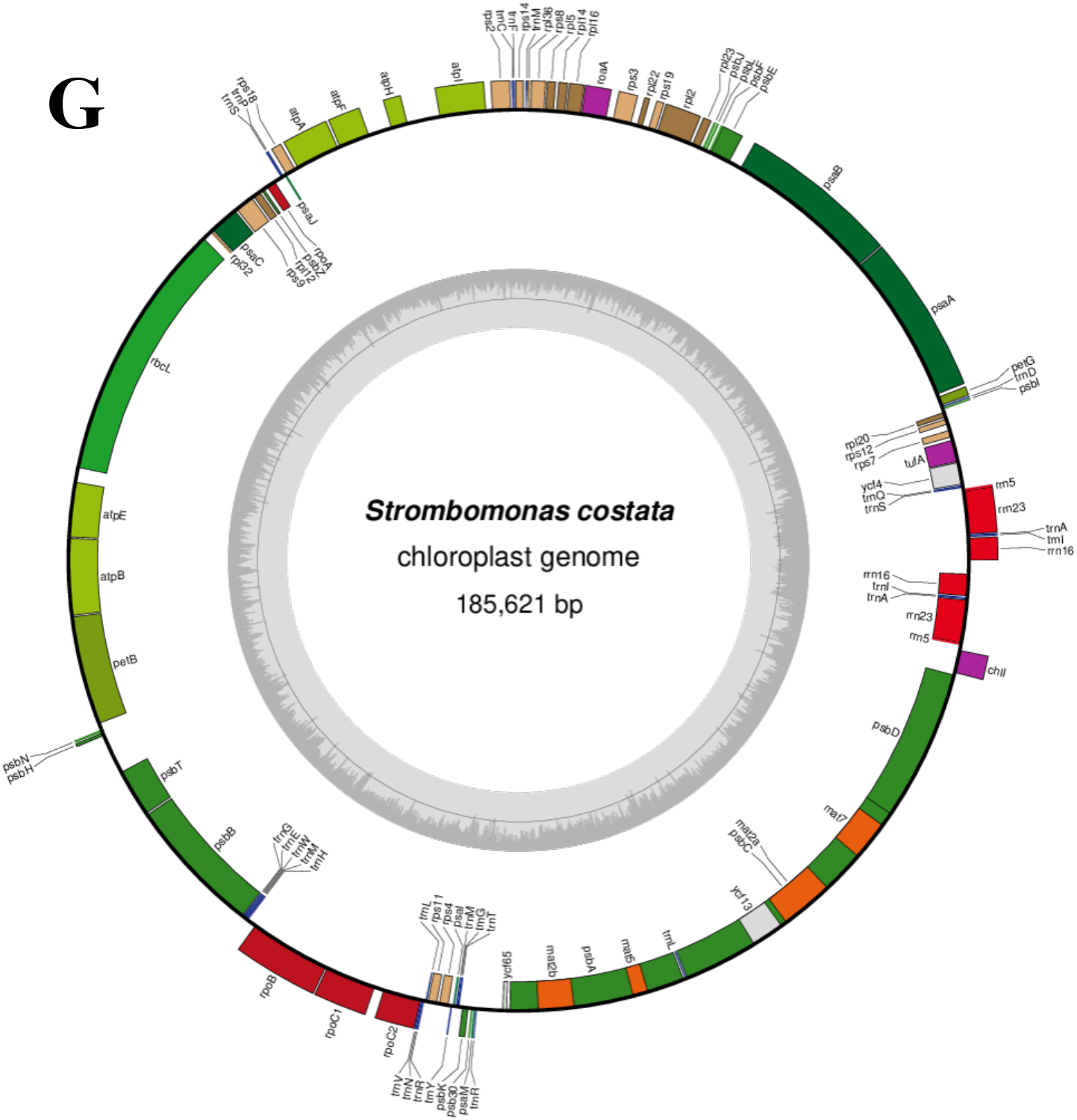
Chloroplast genome maps of *Colacium mucronatum* (A), *Euglena agilis* (B), *Euglena deses* (C), *Euglena undulata* (D), *Flexiglena variabilis* (E), *Phacus warszewiczii* (F) and *Strombomonas costata* (G). Maps were generated using OGDraw v1.3.1 online tool.

